# StainedGlass: Interactive visualization of massive tandem repeat structures with identity heatmaps

**DOI:** 10.1101/2021.08.19.457003

**Authors:** Mitchell R. Vollger, Peter Kerpedjiev, Adam M. Phillippy, Evan E. Eichler

**Affiliations:** Genome Sciences, University of Washington School of Medicine, Seattle, WA, USA; Reservoir Genomics LLC, Oakland, CA; Genome Informatics Section, National Human Genome Research Institute, National Institutes of Health, Bethesda, MD, USA; Howard Hughes Medical Institute, University of Washington, Seattle, WA, USA

## Abstract

**Summary:** Visualization and analysis of genomic repeats is typically accomplished through the use of dot plots; however, the emergence of telomere-to-telomere assemblies with multi-megabase repeats requires new visualization strategies. Here, we introduce StainedGlass which can generate publication quality figures and interactive visualizations that depict the identity and orientation of multi-megabase repeat structures at a genome-wide scale. The tool can rapidly reveal higher-order structures and improve the inference of evolutionary history for some of the most complex regions of genomes.

**Availability and implementation:** StainedGlass is implemented using Snakemake and is available open source under the MIT license at https://mrvollger.github.io/StainedGlass/.

## 1 Introduction

Dot plot analyses are often used to reveal the underlying structure of complex repeats including differences in sequence identity and orientation (Gibbs and McIntyre, 1970; Cabanettes and Klopp, 2018; Rice *et al*., 2000; Sonnhammer and Durbin, 1995). Advances in long-read sequencing technology, however, have recently made more complex repeat structures and genetic variation available for sequence analysis (Chaisson *et al*., 2015; Audano *et al*., 2019; Ebert *et al*., 2021). With increasingly contiguous assemblies of reference genomes (Rhie *et al*., 2021) and complete human chromosomes (Miga *et al*., 2020; Logsdon *et al*., 2021; Nurk *et al*., 2021), complete centromeres, tandem duplications, and other heterochromatic arrays can now be systematically analyzed in their entirety. The size and complexity of these structures, often many megabase pairs in size, elude traditional dot plot analyses for three reasons: 1) current visualization methods are largely based on perfect or k-mer matches which do not lend themselves to the complex higher-order repeats found in centromeres (Willard and Waye, 1987) and the expected mismatches between these large repeats, 2) dot plots offer limited resolution for tandem arrays consisting of megabases of sequence data, frequently reducing them to black squares that relay little information other than the size and presence of a repeat, and 3) the number of possible pairwise matches increases rapidly when identifying exact matches in tandem arrays (e.g. in MUMmer) and this problem is exacerbated further when using a small minimum match length for comparing more divergent arrays.

In this work, we present StainedGlass, which generates identity heatmaps based on sequence alignment rather than small k-mers using an easy, scalable, and customizable workflow that allows for interactive use as well as publication-ready figures. The tool can be applied to study repeat structures at a genome-wide scale or focused at particular regions to characterize complex higher-order repeat structures. As part of our recent analysis of chromosome 8, we developed a prototype of this method which rendered the higher-order repeat structure of the 2 Mbp centromere as an identity heatmap. This prototype facilitated the discovery of higher level symmetry and a layered organization in the centromere, which assisted in the development of a more refined model for centromere evolution, as well as the discovery of hotspots for copy number variation (Logsdon *et al*., 2021).

### 2 Methods

To generate pairwise sequence identity heatmaps for StainedGlass the input sequence is fragmented into windows of a configurable size (default 1 kbp). All possible pairwise alignments between the fragments are calculated using minimap2 (Li, 2018). The color gradient used in the heatmap is then determined by the sequence identities of the alignments which are calculated as: 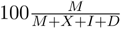 where *M* is the number of matches, *X* the number of mismatches, *I* the number of insertion events, and *D* the number of deletion events. When there are multiple alignments between the same two sequence fragments all alignments other than the one with the most matches are filtered out regardless of their sequence identity. This situation often arises when aligning tandem repeats where there can be multiple valid alignments between fragments, and this strategy assists in highlighting the most representative alignment.

StainedGlass generates two types of outputs: fixed resolution static figures as well as multi-resolution Cooler files (Abdennur and Mirny, 2020) suitable for interactive multiscale visualization using HiGlass (Kerpedjiev *et al*., 2018). Sequence identity in Cooler files is calculated using the method described above for the highest resolution and interpolated by averaging values for lower resolutions. The static figures are more appropriate for visualization of relatively small regions (30 Mbp or less) at publication quality (Figure 1) while the HiGlass visualization is better suited for data exploration of whole genome alignments, such as the sequence identity relationships among the short arms of human acrocentric chromosomes ([interactive HiGlass browser](https://resgen.io/viewer/Ide3NFCTRxWpCgRT2BPFsg)) (Nurk *et al*., 2021; Kerpedjiev *et al*., 2018).

**Figure 1.**
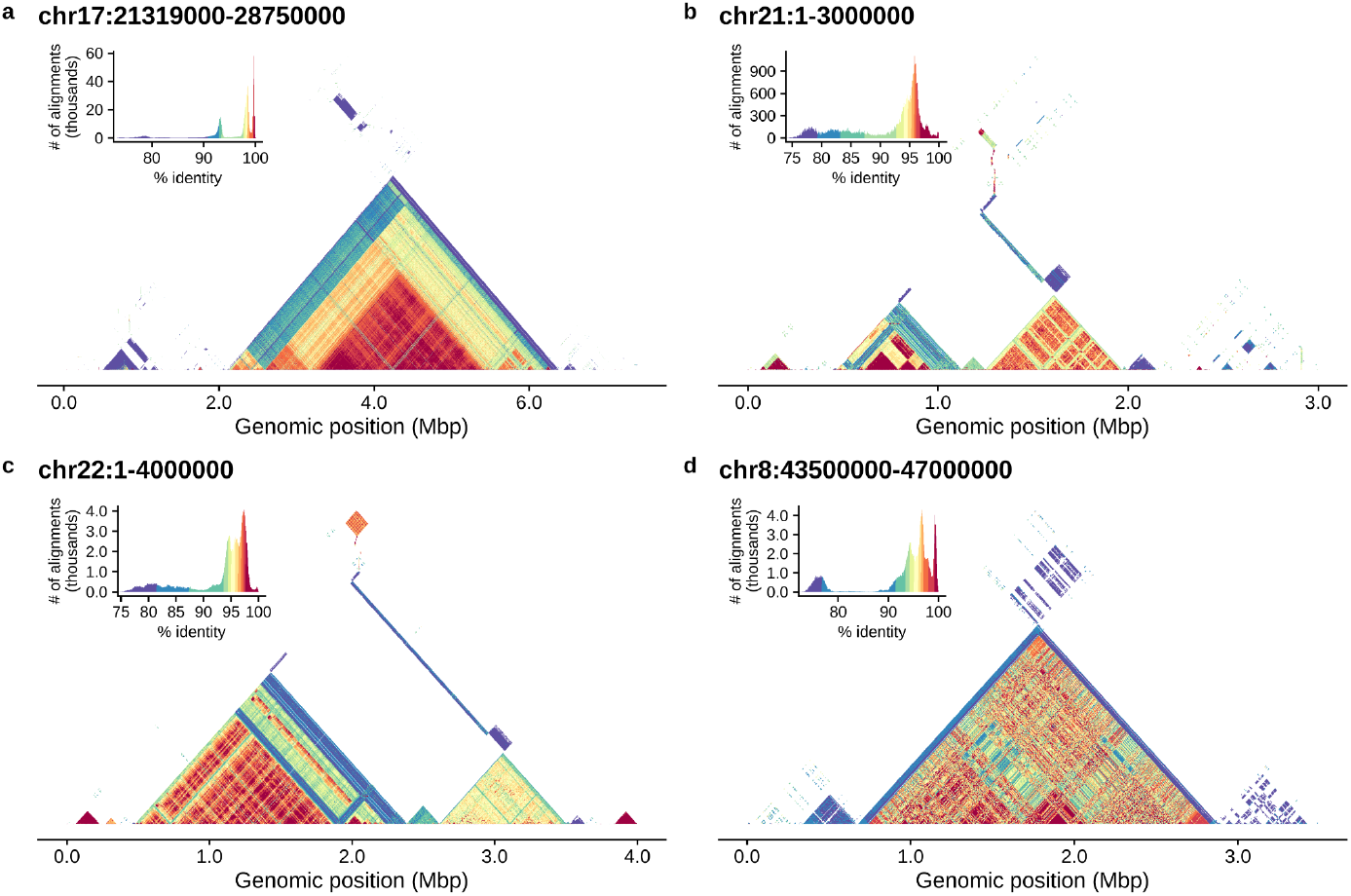
Tandem arrays in T2T-CHM13 visualized with StainedGlass. This figure was generated entirely by StainedGlass and has not been edited or refined in any way. a) The chromosome 17 centromere dot plot reveals several different higher order repeat arrays which are made apparent by the colored percent identity. b,c) Show the beginning of the acrocentric short arms of chromosomes 21 and 22. d) Highlights the centromeric array on chromosome 8 first presented by (Logsdon *et al*., 2021). An interactive browser of this data is available at resgen.io.

The tool is made available using Snakemake (Köster and Rahmann, 2012, 2018; Mölder *et al*., 2021) which allows for reproducible and scalable data analyses. The stability of new changes is automatically tested with each new addition using continuous integration via GitHub actions. Finally, StainedGlass is a Snakemake standard compliant workflow and therefore has automated [usage documentation](https://snakemake.github.io/snakemake-workflow-catalog/?usage=mrvollger%2FStainedGlass) on the Snakemake website.

### 3 Usage and examples

StainedGlass is run using the Snakemake workflow language. To run StainedGlass, clone the repository from https://github.com/mrvollger/StainedGlass and follow the installation instructions. From the cloned directory you can then generate the alignments for StainedGlass using the following command.

~~~
       snakemake --use-conda --cores 24 \
           --config fasta={path.to.fasta} sample={sample.prefix.for.output}
~~~

All configuration options are described in `config/README.md` and parameters can be specified on the command line as done above or by modifying the configuration file (`config/config.yaml`). You can also preview the jobs that will be run without executing the pipeline by adding `-n` to the command line. To generate static identity heatmaps from the StainedGlass alignments, add `make_figures` to the Snakemake command. For an example of the output of this command see Figure 1.

~~~
      snakemake --use-conda --cores 24 \
           --config fasta={path.to.fasta} sample={sample.prefix.for.output} \
           make_figures
~~~

StainedGlass can also be used to make Cooler (Abdennur and Mirny, 2020) files that can be loaded into HiGlass for whole genome visualizations. An example of this interactive browser for the telomere to telomere assembly of CHM13 can be found at resgen.io (Nurk *et al*., 2021)

~~~
      snakemake --use-conda --cores 24 \
              --config fasta={path.to.fasta} sample={sample.prefix.for.output} \
              cooler
~~~

The resulting Cooler files can then be easily visualized on a local browser using HiGlass.

~~~
      pip install higlass-manage
      higlass-manage view results/{sample}.{\d+}.{\d+}.mcool
~~~

## 4 Conclusion

We developed StainedGlass, a visualization tool for large genomic repeats that displays the higher-order structure and sequence relationship of these complex regions. The underlying Snakemake workflow makes StainedGlass both reproducible and scalable at the whole genome level. StainedGlass produces static output visualizations which are publication ready, and has additional options for interactive data exploration with HiGlass.

## Acknowledgements

The authors thank April Lo and Michelle Noyes for help in editing this manuscript and G. A. Logsdon for aesthetic suggestions.

## References

Abdennur, N. and Mirny, L.A. (2020) Cooler: scalable storage for Hi-C data and other genomically labeled arrays. Bioinformatics, 36, 311–316.

Audano, P.A. et al. (2019) Characterizing the Major Structural Variant Alleles of the Human Genome. Cell, 176, 663–675.e19.

Cabanettes, F. and Klopp, C. (2018) D-GENIES: dot plot large genomes in an interactive, efficient and simple way. PeerJ, 6, e4958.

Chaisson, M.J.P. et al. (2015) Resolving the complexity of the human genome using single-molecule sequencing. Nature, 517, 608–611.

Ebert, P. et al. (2021) Haplotype-resolved diverse human genomes and integrated analysis of structural variation. Science.

Gibbs, A.J. and McIntyre, G.A. (1970) The diagram, a method for comparing sequences. Its use with amino acid and nucleotide sequences. Eur. J. Biochem., 16, 1–11.

Kerpedjiev, P. et al. (2018) HiGlass: web-based visual exploration and analysis of genome interaction maps. Genome Biol., 19, 125.

Köster, J. and Rahmann, S. (2012) Snakemake—a scalable bioinformatics workflow engine. Bioinformatics, 28, 2520–2522.

Köster, J. and Rahmann, S. (2018) Snakemake-a scalable bioinformatics workflow engine. Bioinformatics, 34, 3600.

Li, H. (2018) Minimap2: pairwise alignment for nucleotide sequences. Bioinformatics, 34, 3094–3100.

Logsdon, G.A. et al. (2021) The structure, function and evolution of a complete human chromosome 8. Nature, 593, 101–107.

Miga, K.H. et al. (2020) Telomere-to-telomere assembly of a complete human X chromosome. Nature, 585, 79–84.

Mölder, F. et al. (2021) Sustainable data analysis with Snakemake. F1000Res., 10, 33.

Nurk, S. et al. (2021) The complete sequence of a human genome. bioRxiv, 2021.05.26.445798.

Rhie, A. et al. (2021) Towards complete and error-free genome assemblies of all vertebrate species. Nature, 592, 737–746.

Rice, P. et al. (2000) EMBOSS: the European Molecular Biology Open Software Suite. Trends Genet., 16, 276–277.

Sonnhammer, E.L.L. and Durbin, R. (1995) A dot-matrix program with dynamic threshold control suited for genomic DNA and protein sequence analysis. Gene, 167, GC1–GC10.

Willard, H.F. and Waye, J.S. (1987) Hierarchical order in chromosome-specific human alpha satellite DNA. Trends Genet., 3, 192–198.

